# A shared neural code for gender across faces, bodies, and objects in the human brain

**DOI:** 10.64898/2026.05.04.722201

**Authors:** Wenjie Liu, Xiqian Lu, Yuhui Cheng, Ruidi Wang, Xinquan Lu, Xiangyong Yuan, Yi Jiang

## Abstract

Humans can readily infer gender from diverse visual categories, including faces, bodies, and objects, yet it remains unclear whether the brain constructs a shared, category-general representation of gender. To fill the gap, we measured neural responses using fMRI while participants viewed female and male stimuli across these three categories. Multivoxel pattern analyses revealed that gender information was encoded throughout distributed occipitotemporal regions for all categories. Critically, cross–category decoding and regression–based representational similarity analyses converged on a region of the right middle temporal gyrus (rMTG) that encoded gender independently of stimulus category. This region also carried category information, indicating mixed selectivity. Comparisons between neural representational dissimilarity matrices and those from a fine-tuned convolutional neural network (CNN) showed that rMTG activity most closely resembled intermediate CNN layers, suggesting that shared gender representations rely on mid-level visual features. Finally, functional connectivity analyses revealed that gender-related interactions among occipitotemporal regions were similar for faces and bodies but not for objects. Together, these findings identify a category-general neural hub for gender perception and illuminate how mid-level visual features and distributed networks support the abstraction of gender across heterogeneous visual inputs.

## 1 Introduction

The exceptional ability to recognize gender is vital for mate choice in many species. It is also critical for social interactions in our daily life, as a precise gender perception often guides appropriate behaviors in the interaction. With an influx of visual signals, gender can be inferred from a variety of sources, each with notably different characteristics (Brown & Perrett, 1993; Gandolfo & Downing, 2020; Lemm et al., 2005; Webster et al., 2004). For instance, our visual system can readily determine gender from faces, even when the face stimuli are partially degraded, occluded or silhouetted (Cellerino et al., 2004; Davidenko, 2007; Dupuis-Roy et al., 2009). Bodies and objects also provide cues for recognizing gender in real-world visual contexts (Gandolfo & Downing, 2020; Ghuman et al., 2010; Javadi & Wee, 2012; Lemm et al., 2005). Thus, our representation of gender is independent of specific input cues, suggesting that gender may be encoded in a relatively abstract manner. This prompts an important question of whether there are shared neural correlates for gender perception from faces, bodies and objects.

Most previous studies on the neural basis of gender perception have primarily focused on faces, since faces are a key source of gender cues. Those studies have identified a wide range of brain areas involved in processing face gender, including occipital brain regions, fusiform gyrus, insula, inferior frontal gyrus, orbitofrontal cortex (Contreras et al., 2013; Freeman et al., 2010; Kaul et al., 2011). Despite that bodies and objects also dedicate to gender perception (Gandolfo & Downing, 2020; Ghuman et al., 2010; Javadi & Wee, 2012; Lemm et al., 2005), the neural basis of gender perception from the two categories has received much less attention. Only one neuroimaging study using static body images revealed widespread brain regions for body gender, including large parts of the occipitotemporal cortex and clusters in parietal and frontal regions (Foster et al., 2019), a pattern similar to those reported for facial gender processing. Therefore, it remains unclear not only how the brain processes object gender, but also whether there are gender-specific regions that encode gender across different categories. Furthermore, given that gender is extracted from distinct categories, do the gender-specific regions overlap with category-selective regions, or occupy anatomically separate regions? Elucidating this relationship would provide a comprehensive understanding of the mechanisms underlying gender processing.

To investigate these issues, the current study employed functional magnetic resonance imaging (fMRI) to record participants’ brain activity while they viewed images of bodies, and objects varying in gender-related cues. First, we conducted a whole-brain multivoxel pattern analysis (MVPA) on neural responses evoked by female and male stimuli across three categories (faces, bodies and objects), to identify brain regions exhibiting within-category gender representations and their potential overlap that may imply shared gender encoding. In parallel, we performed a cross-category MVPA (e.g., trained on faces, tested on bodies) to confirm shared gender representations in the brain region capable of encoding gender across categories. The resulting brain map for gender representations was compared with those for category representations to visualize the degree of overlap between these two coding schemes. Second, we applied a searchlight representational similarity analysis (RSA) for the whole brain and predicted the neural representation dissimilarity matrices (RDMs) with gender and category RDMs simultaneously. This disentangled gender and category representations and provided converging evidence for the shared gender-encoding region identified by MVPAs. The neural RDMs in the region were further compared with gender RDMs computed at different layers of a convolutional neural network (CNN) to characterize the hierarchical nature of the visual features encoded by these brain regions. Finally, we conducted functional connectivity analyses for all gender-related regions—combination of all regions that encode gender from faces, bodies and objects—to elucidate the similarity of the neural network in processing gender from different categories. In summary, by analyzing neural response patterns and inter-area functional connectivity, this study comprehensively characterizes how the brain constructs generalized gender representations from distinct visual inputs, bridging category-specific and category-general mechanisms of gender perception.

## 2. Results

### 2.1 Whole-brain classification of gender within category

The whole-brain searchlight MVPA within the three categories (Fig. 1a) delineated the brain regions underlying the gender representation of face, body, and object, respectively. For face stimuli, the MVPA results (Fig. 2a, left panel) showed significant above-chance classification of gender in bilateral cuneus (CUN), calcarine (CAL), lingual gyrus (LG), superior occipital gyrus (SOG), middle occipital gyrus (MOG), middle temporal gyrus (MTG); right superior parietal lobule (SPL); left fusiform gyrus (FG). For body stimuli, classification of gender (Fig. 2a, middle panel) was significantly above chance in a large portion of occipitotemporal cortex, and clusters in bilateral SPL, IPL, postcentral gyrus (PoCG). For object stimuli, the gender was classified in bilateral CUN, CAL, LG, FG, SOG, MOG, IOG, ITG; right MTG (Fig. 2a, right panel). These results show that distributed brain regions were sensitive to gender encoding of faces, bodies, and objects, mainly in the occipitotemporal cortex. More importantly, we examined the anatomical overlap across these regions to provide potential clues for the neural substrates underlying category-general gender representation. As illustrated in Fig. 2c (yellow areas), the brain regions able to decode gender in all three categories (faces, bodies, and objects) included bilateral CUN, CAL, LG, SOG, MOG; left FG, IOG; and right MTG.

**Fig. 1.**
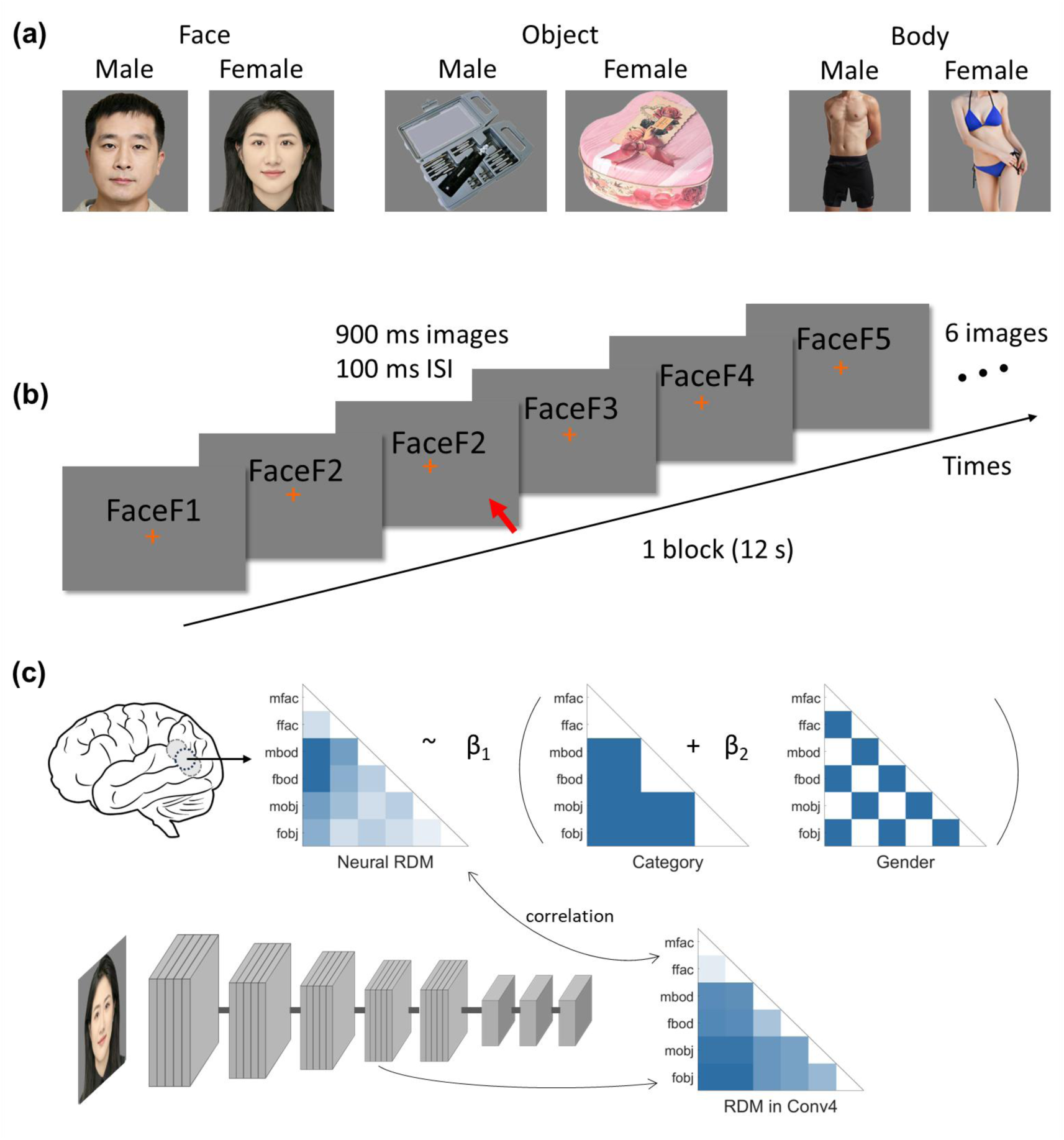
Schematic diagram of stimuli and procedure in fMRI experiment. (a) Schematic diagram of stimuli for the 6 conditions in the fMRI experiment. The images from left to right are example for images of male face, female face, male object, female object, male body and female body, respectively. (b) Schematic diagram of a block in fMRI experiment. Participants viewed the stimuli in 12 s blocks, where each block contained images from one condition (female faces shown here). Each image with fixation was presented for 900 ms, followed by a 100 ms fixation between images. The red arrow indicates a repeated image, where the participant should press button. (c) Top panel: Searchlight regression-based RSA was conducted to predict neural RDM structure using a linear combination of theoretical category and gender model RDMs within each searchlight sphere. Bottom panel: RSA linking activation in brain and CNN. Correlations were calculated between RDMs from each layer of a fine-tuned AlexNet and neural RDMs from each region of interest. The example illustrates the correlation calculation between Conv4 layer RDM and neural RDM. For privacy reasons, the example faces are the authors’ own.

**Figure 2.**
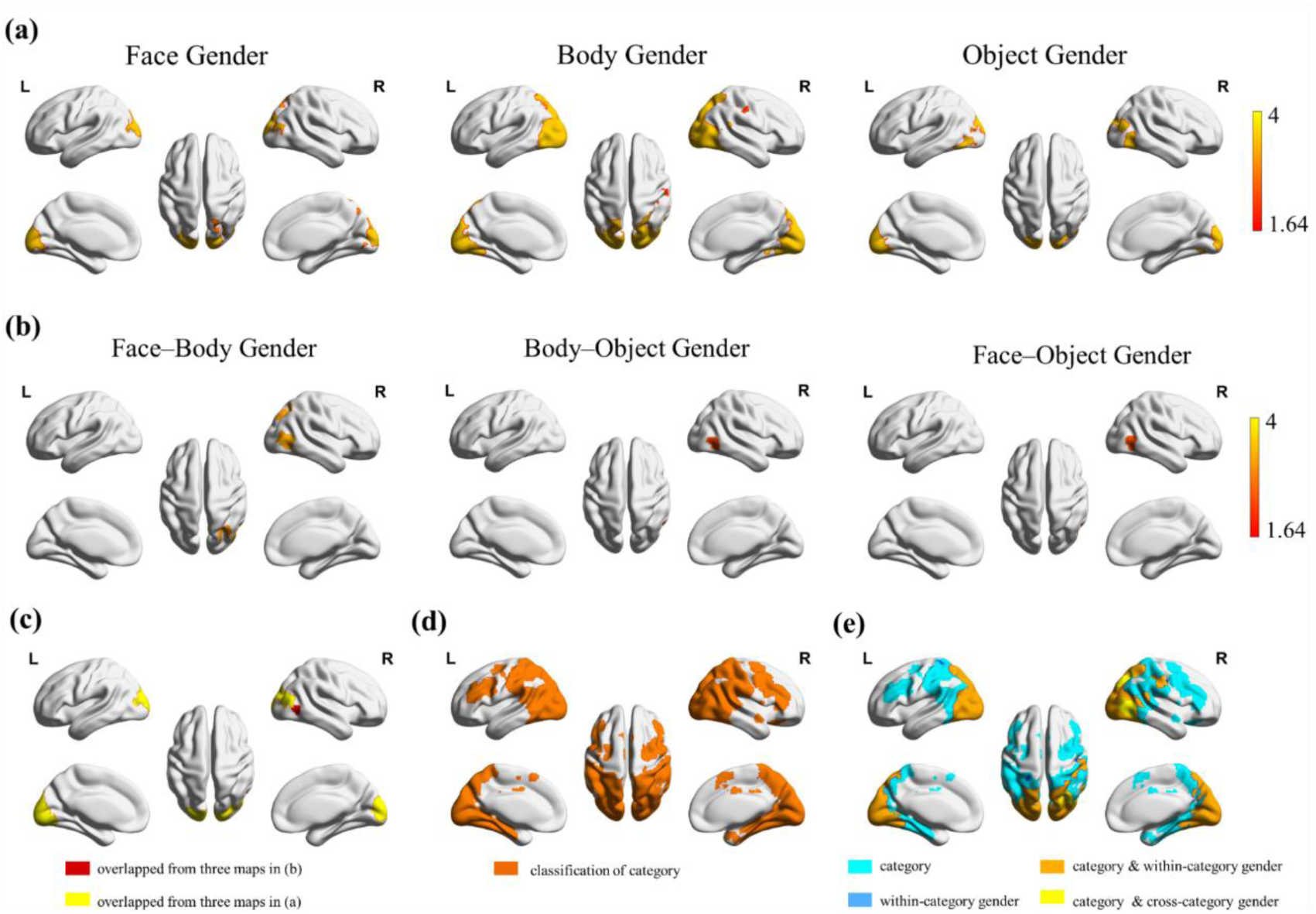
Results of whole-brain searchlight analysis for the classification of gender and category. (a) Brain regions able to classify gender from faces (left panel), bodies (middle panel), and objects (right panel). (b) Brain regions able to classify gender across faces and bodies (left panel), across bodies and objects (middle panel), and across objects and faces (right panel). The statistical maps in (a) and (b) show significant brain regions after Monte Carlo simulations (*p* = 0.001 at the initial voxel-wise, 5000 iterations; the color scale bar show the z-scores above 1.64, corresponding cluster level *p* < 0.05). (c) The yellow color depicting overlap areas of three maps for within-category gender classification in (a). The red color depicting overlap areas of three maps (rMTG) for three cross-category gender classification maps (b). (d) Brain regions for category classification, identified by the conjunction analysis of significant maps of category classification on each pairwise categories. (e) The cyan color shows the regions able to classify category. The orange and yellow show the overlapping regions for category classification and within- category, cross-category gender classification, respectively. The blue color shows a few regions able to classify within-category gender, but not category.

### 2.2 Whole-brain classification of gender across categories

To provide more compelling evidence for category-general neural representation of gender, we conducted a cross-category classification of gender using whole-brain searchlight analysis. Specifically, we trained classifier to distinguish gender using neural activity from one category (e.g., face), and then tested the classifier on neural activity from another category (e.g., body), and vice versa. The results were showed in Figure 2b. Across faces and bodies (Fig. 2b, left panel), the cross-category gender classification was significantly above chance in right MOG, MTG, ITG, FG, SOG, SPL, IPL, angular. Across bodies and objects (Fig. 2b, middle panel), the cross-classification of gender was significantly above chance in right MOG, MTG, ITG. Across objects and faces (Fig. 2b, right panel), the cross-classification of gender was significantly above chance in right MOG, MTG, ITG.

A conjunction analysis showed that the three significant cross-classification maps overlapped in a cluster of right MTG (rMTG, MNI coordinates: 48, −62, 0, including 73 voxels), as colored by red in Figure 2c. In other words, the cluster in rMTG could significantly classify neural activity patterns of gender across the three categories, indicating a shared neural representation of gender. Notably, the rMTG identified here is anterior and contiguous to the rMTG identified by the conjunction analysis of the within-category MVPA (Fig. 2c, the yellow clusters neighboring to the red clusters). This finding collectively indicated that the rMTG plays a critical role in category-general gender encoding.

### 2.3 Whole-brain classification of category

To determine whether the identified gender-related brain areas specifically encode gender or they could also encode category, we conducted another whole-brain searchlight MVPA on category classification. The brain regions representing category were defined as those where neural patterns significantly differed between faces and bodies, faces and objects, and bodies and objects. These brain regions (Fig. 2d) included much of occipital, posterior, and parietal lobe, and clusters in bilateral middle frontal gyrus (MFG) and inferior frontal gyrus (IFG). The conjunction analysis revealed the overlapping regions of gender representation and category representation. As shown in Fig. 2e, most gender-encoding regions, including within-category (orange color) and cross-category (yellow color) gender representation, are also able to classify category. A few exceptions (blue color) that encoded within-category gender but could not classify category included small clusters (all < 50 voxels) in CUN, CAL, LG, FG, IOG, MOG, PoCG. The findings indicates that gender representation does not fully separate from category representation and implicate completely distinct brain regions, reflecting mixed selectivity of the neural responses in these regions.

### 2.4 Shared gender representation independent of category selectivity

Considering the gender representation are mixed with category representation across brain, we conducted a regression-based searchlight RSA on the whole brain to isolate and examine gender-specific representation independent of category selectivity (Fig. 1c). As shown in Fig. 3a, category RDM significantly predicted neural RDM in a wide range of brain regions, consistent with the whole-brain classification results (Fig. 2d). More importantly, gender RDM significantly predicted neural RDMs in a cluster (MNI coordinates: 47, −61, 1, 172 voxels) mainly located in rMTG (137 voxels) and rITG (23 voxels). The location of the cluster aligned with the rMTG identified by cross-category MVPA, strongly supporting the critical role of rMTG in representing gender even with the category representation excluded.

**Figure 3.**
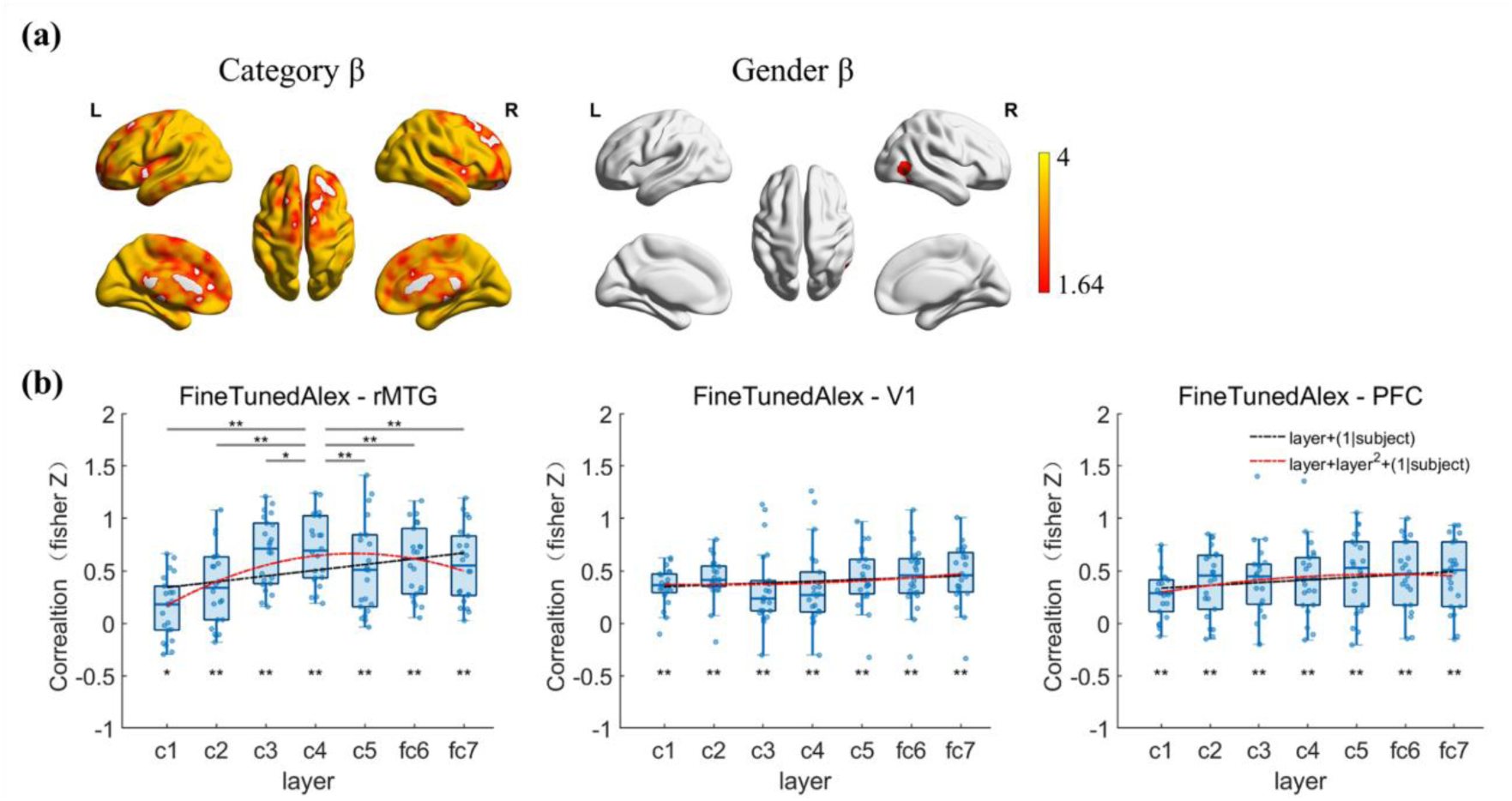
Results of RSA. (a) Regression-based RSA results depicting areas whose activation were significant predicted by category and gender theoretical RDMs. These statistical maps show the significant brain regions after Monte Carlo simulations (*p* = 0.001 at the initial voxel-wise, 5000 iterations; the color scale bar show the z-scores above 1.64, corresponding cluster level *p* < 0.05). (b) RSA linking neural responses in rMTG (left panel), V1(middle panel) and PFC (right panel) to CNN layers. * *p* < 0.05, ** *p* < 0.01 (FDR- corrected). The data predicted by the linear and quadratic model were plotted using red and black dashed lines.

### 2.5 Hierarchical nature for shared gender representation

The evidence from MVPA and RSA consistently showed that rMTG encoded gender across categories. To determine which level of features drives this shared gender representation, we computed the similarity between neural RDMs in rMTG and the CNN RDMs in seven representative layers of a fine-tuned gender-classification AlexNet (Fig. 1c). To directly compare with brain regions processing low-level and high-level features, we also examined the similarity between neural RDMs in V1 and PFC and the CNN RDMs across layers. The analysis revealed that all the neural RDMs in the three brain regions are significantly correlated with the CNN RDM in each layer (*ps* < 0.05). However, a quadratic trend of the correlation across layers was most robustly found in rMTG (linear 𝑅 ^2^𝑎𝑑𝑗 = 0.75, quadratic 𝑅 ^2^𝑎𝑑𝑗 = 0.86; linear BIC = 25.46, quadratic BIC = −39.58), whereas the linear and the quadratic trend fit comparably well in V1 (two models’ 𝑅 ^2^were 0.71; linear BIC = −44.42, quadratic BIC = −40.66) and PFC (linear 𝑅 ^2^𝑎𝑑𝑗 = 0.84, quadratic 𝑅 ^2^𝑎𝑑𝑗 = 0.85; linear BIC = −81.57, quadratic BIC = −83.27). These results confirmed our hypothesis: intermediate-level gender-related features are represented in rMTG, in contrast to V1 and PFC. To identify the optimal layer best matching the neural activity in rMTG, we sorted the correlation coefficients along the layers. The strongest correlation was found at layer 4 (i.e., Conv4), and significantly larger than those at other layers (*ps* < 0.05, Fig. 3b left panel, Table S1). These findings were replicated using VGG16 (see details in Supporting Information), further showing the generalizability of the findings across models. Together, these findings indicate that shared gender representation in rMTG relies on mid-level features, naturally distinct to low-level or high-level representations in V1 and PFC.

### 2.6 Shared neural network for gender representation across categories

In addition to the shared neural hubs for gender representation, we also investigated the functional connectivity underlying gender representation to uncover the commonalities in inter-area transmission of gender-related information across categories. Using PPI analysis, we first examined whether functional connectivity between gender-related regions — those able to decode gender of face, body and object—exhibited significant differences for male versus female stimuli within each category (Fig. 4). The results showed that when processing male versus female faces, no significant differences in inter-regional functional connectivity was observed (Fig. 4a); however, when processing male versus female bodies, the functional connectivity significantly differed mainly between CAL, CUN, LING and SOG, MOG, IOG, FG (Fig. 4b, see details in Table 1), and when processing male versus female objects, significantly different functional connectivity was found between left CAL and bilateral FG, between left LING and bilateral FG (Fig. 4c). Above all, these findings suggest that gender-related information transmission was mainly occurs between regions located in the posterior occipitotemporal areas.

**Figure 4.**
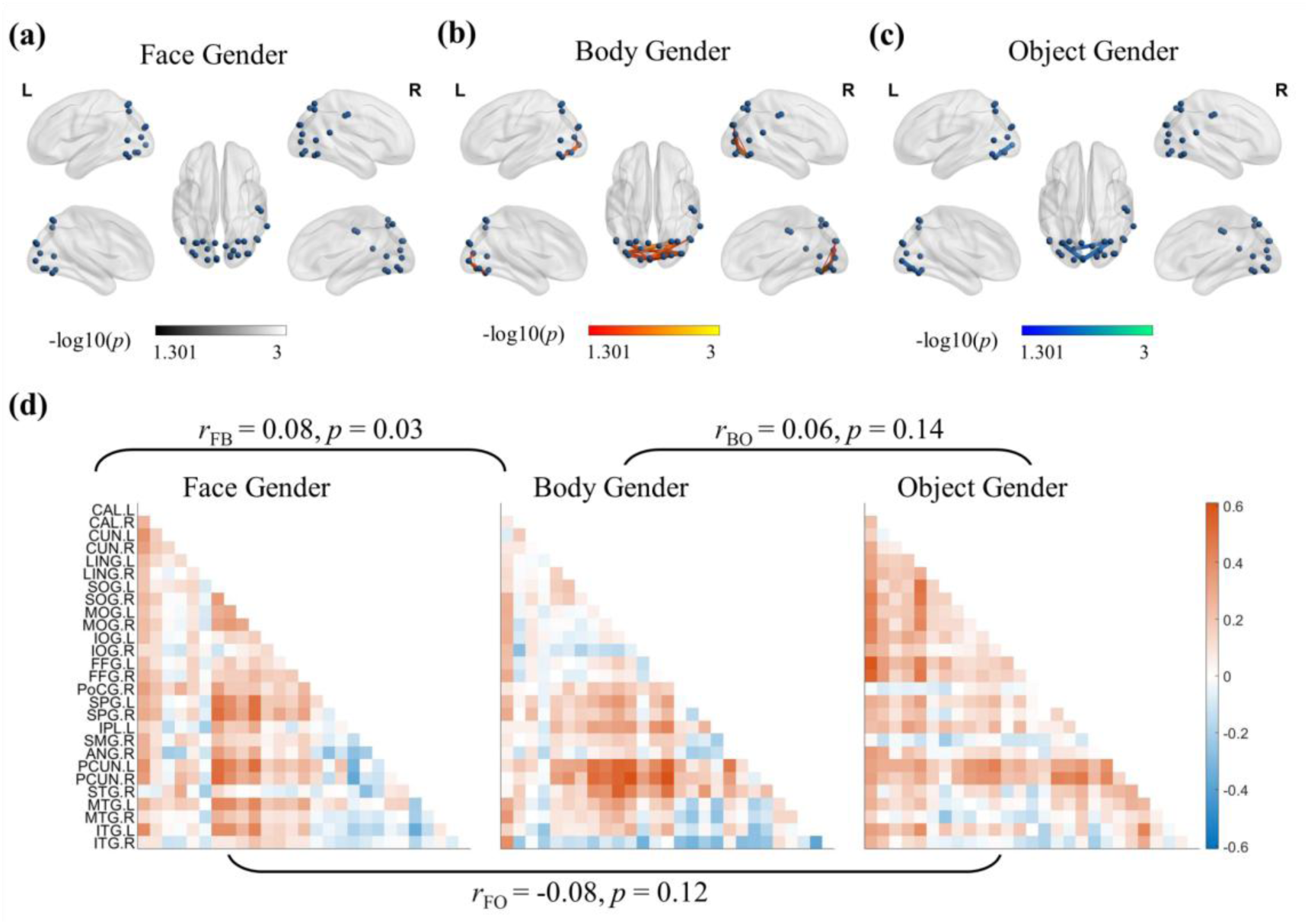
Results of the PPI analysis. (a) – (c) The significant functional connectivity (lines in each figure) changes between seeds (blue dots) for the gender processing of face (a), body (b) and object (c). The color scale bar shows −log_10_(*p* values) ranging between 1.301 (*p* =0.05) and 3 (*p* =0.001), FDR-corrected. (d) PPI matrix for gender processing of face, body and object, and the correlation between these marix. Each PPI value in matrix showed in figure was the mean PPI from all participants.

**Table 1.**
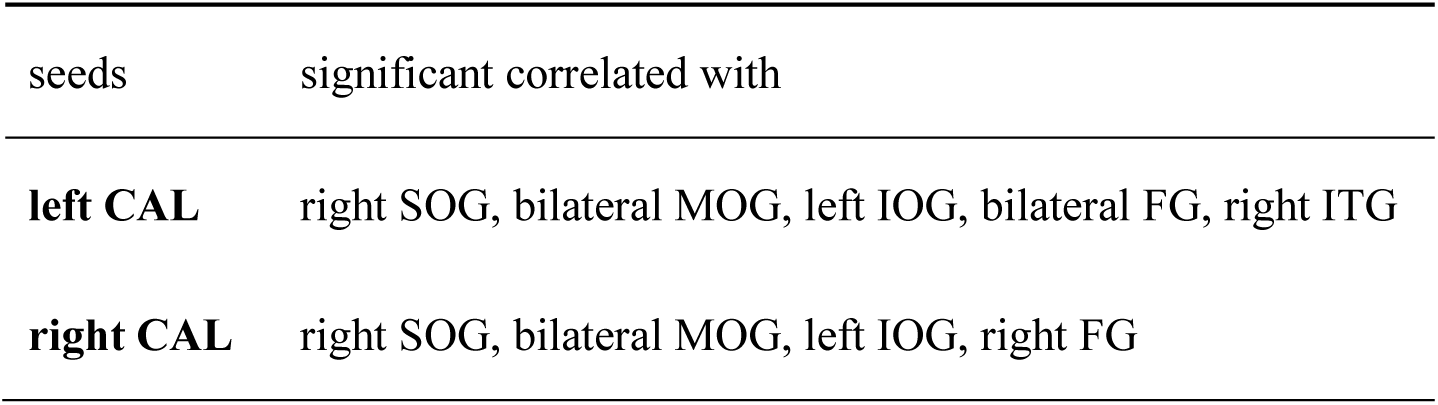
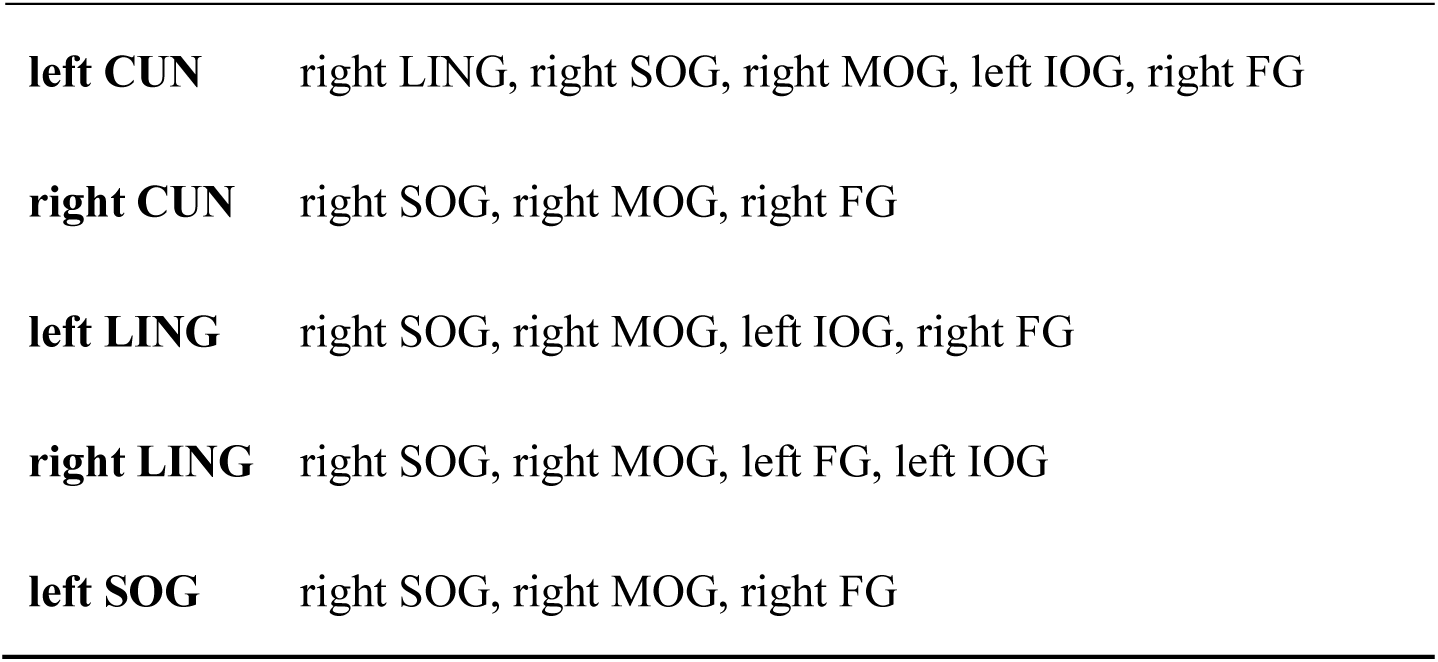
Significant PPI differences when processing male versus female bodies.

| seeds | significant correlated with |
| --- | --- |
| **left CAL** | right SOG, bilateral MOG, left IOG, bilateral FG, right ITG |
| **right CAL** | right SOG, bilateral MOG, left IOG, right FG |
| **left CUN** | right LING, right SOG, right MOG, left IOG, right FG |
| **right CUN** | right SOG, right MOG, right FG |
| **left LING** | right SOG, right MOG, left IOG, right FG |
| **right LING** | right SOG, right MOG, left FG, left IOG |
| **left SOG** | right SOG, right MOG, right FG |

Next, we calculated the correlation between the PPI value matrix of the pair-wised brain regions to quantify the similarity of the functional connectivity in gender processing across categories (Fig. 4d). The results showed that functional connectivity patterns of face and body gender processing were significantly correlated (*r*_FB_ = 0.08, *p* = 0.03). However, functional connectivity patterns showed no significant relationship between body and object (*r*_BO_ = 0.06, *p* = 0.14) or between face and object (*r*_FO_ = −0.08, *p* = 0.12), with the latter showing a negative trend. In summary, these findings suggest that the functional connectivity of gender processing is similar for faces and bodies, revealing shared neural networks for gender representation in these two categories.

## 3 Discussion

Our study revealed the neural substrates underpinning gender perception from facies, bodies and objects. Consistent with prior findings (Foster et al., 2019; Freeman et al., 2010; Kaul et al., 2011; Wang et al., 2023), face or body gender could be encoded in distributed brain regions, mainly in the occipitotemporal lobe. The encoding of object gender in our study was observed in similar regions. More importantly, among those gender-related regions, the rMTG was able to classify gender across the three categories, indicating a category-general, shared neural representation. Moreover, these gender- related regions could also classify category, indicating mixed-selectivity of neural responses in these regions. Furthermore, the neural responses in rMTG encode shared gender most similarly to the artificial activities in the intermediate layer of CNN, revealing that shared gender representation relied on mid-level visual features. In addition, the neural network for face and body gender processing also exhibited a significant similarity in functional connectivity. These findings reveal shared neural mechanisms for category-general gender perception: the rMTG, serving as a critical local hub with mixed-selectivity and relying on mid-level features, along with functionally similar distributed networks for faces and bodies.

### 3.1 Neural correlates of gender perception from facies, bodies and objects

Numerous previous findings show that face gender processing activates extensively distributed brain region, including OFA, FFA, lateral FG etc. as the core brain regions for face processing (Contreras et al., 2013; Freeman et al., 2010; Kaul et al., 2011; Wiese et al., 2012), and insula, inferior frontal gyrus etc. as the peripheral brain regions for face processing (Freeman et al., 2010; Kaul et al., 2011). Consistently, we also found a distributed brain network for encoding face gender, including bilateral CUN, CAL, LG, SOG, MOG, MTG, right SPL, and left FG. Searchlight analysis also revealed that body gender could be represented in extensive brain areas, including much of occipitotemporal cortex, bilateral SPL, IPL, PoCG. Echoed with our findings, prior studies using static bodies (Foster et al., 2019) and biological motion (Wang et al., 2023) also revealed distributed brain areas sensitive to body gender, such as clusters in occipitotemporal and parietal lobe. As for objects, the current study was the first to explore its gender encoding local hubs, revealing key regions including bilateral CUN, CAL, LG, FG, SOG, MOG, IOG, ITG and right MTG. In summary, the within-category classification showed distributed brain regions for gender representation of facies, bodies or objects.

Apparently, body gender processing engaged a more extensive distributed network compared to face or object gender processing. It is possible that gender information may be more salient or prioritized in body processing, as previous findings showed that body gender processing was resistant to interference from body weight (Johnstone & Downing, 2017), and involves wider brain regions than weight-related processing (Foster et al., 2019). In contrast, face and object gender processing may involve greater competition with other stimulus dimensions (e.g., face identity) , potentially attenuating gender-related neural signals, making its neural signature more difficult to isolate.

Those distributed regions were not isolated; rather, gender perception also relies on hierarchical processing across cortical levels (Bernstein et al., 2018; Joassin et al., 2011). For instance, face gender processing was linked to significant functional synchronization between FFA and OFA (Bernstein et al., 2018). Moreover, the fusiform gyrus and occipital areas encoded the linear changes of objective face gender, while the subjective perception of face gender were encoded in the orbitofrontal cortex (Freeman et al., 2010). This indicates that face gender encoding proceeds hierarchically, from early areas in fusiform gyrus and occipital to higher-order orbitofrontal cortex. Consistently, the current study was first to reveal transmission underlying body and object gender: significant functional connectivity between upstream regions (e.g., CAL) and downstream areas (e.g., FG) in posterior occipitotemporal areas. However, no significant connectivity for face gender was observed, possibly because gender-irrelevant one-back task weakened gender-related response. This speculation can be supported by previous evidences showing that task relevance enhances neural responses to corresponding facial information processing (Dobs et al., 2018; Foster et al., 2021) .

### 3.2 Shared representation of gender across facies, bodies and objects

Previous studies using adaptation paradigm showed that prolonged exposure to body induced gender aftereffect in face perception, and vice versa (Ghuman et al., 2010; Kessler et al., 2013; Palumbo et al., 2015). Such cross-category adaptation suggests shared gender representation across person-related categories. The current study provides the first direct neuroimaging evidence: gender could be decoded across neural responses to faces and bodies in right MOG, MTG, ITG, FG, SOG, SPL, IPL, angular, suggesting that these regions contribute to the shared representation of gender, particularly for person-related stimuli. However, Foster et al. (2019) did not revealed such shared gender representation. This discrepancy may stem from stimulus differences. Foster et al. (2019) adopted the face without external features, which are vital cues for gender recognition (Rossion, 2002). This manipulation might diminish the perceived gender distinction and discriminability of neural response. In contrast, our ecological face stimuli provided richer gender cues, facilitating the detection of cross-category gender decoding. Moreover, cross-category gender decoding also generalized to objects: right MOG, MTG and ITG can decoded gender across neural responses to objects and faces or objects and bodies, consistent with behavioral evidence that cross-category gender adaptation can extend to non-person stimuli (Javadi & Wee, 2012).

More importantly, the current study examined whether this shared representation extends to all three categories in a more general manner. MVPA and RSA consistently showed that rMTG was a critical local hub for category-general gender representation among faces, bodies and objects. Does this category-general gender representation reflect abstraction from gender-related signals, or merely the representation of low-level visual features shared by distinct categories? Leveraging the established correspondence between CNN layers and the human visual hierarchy (Eickenberg et al., 2017; Güçlü & van Gerven, 2017), our study was the first to reveal that rMTG exhibited a mid-level abstraction for shared gender representation. Similarly, previous studies comparing brain and CNN models revealed abstract representations in temporal lobe for object discrimination (Cichy et al., 2016), and face-selective regions (e.g., inferior partial cortex) for face emotion processing (Zhang et al., 2023). Moreover, it has been substantiated that the temporal lobe contributes to the semantic processing of social information (Binney & Ramsey, 2020; Ralph et al., 2017), and the middle temporal gyrus engages in the integration of audiovisual gender (Casado-Aranda et al., 2018; Li et al., 2015). Taken together, the cross-category gender representation in rMTG is unlikely to arise from low-level visual features processing, but more likely reflects a certain degree of integration of mid-level visual features.

Notably, the identified cluster in posterior rMTG does not exclude other possible downstream areas in the visual processing pathway that encode gender with a degree of more abstraction. The unsuccessful observation for such downstream may be related to the gender-irrelevant task. We speculate that the anterior temporal lobe (ATL) may be the key downstream area, given its role in representing abstract and conceptual social information (Rosler & Amodio, 2022), and its involvement in gender processing (Platonov et al., 2019; Tiesinga et al., 2023). Thus, shared gender representation may involve a distributed brain network: posterior rMTG for mid-level abstraction, subsequently projecting to downstream regions like ATL for higher-level abstraction. This aligns with the third visual pathway proposed in a recent review, in which social information processing extends from the primary visual cortex to the MT, then to the posterior superior temporal sulcus (pSTS), and finally to the anterior superior temporal sulcus (aSTS) (Pitcher & Ungerleider, 2021). Moreover, distributed brain network model for the integration of faces and bodies also presented that the cross-category shared brain regions for a particular dimension (such as identity or emotion) involve lower-level occipitotemporal lobe and higher-level STS and ATL (Foster, 2022; Foster et al., 2021; Peelen et al., 2010). This model also suggests a close link between face and body processing as person-related stimuli, potentially explaining why similar functional connectivity was only observed for face and body gender. However, the above proposed mechanisms for shared gender representation, including the specific contribution of ATL and the neural transmission between rMTG and ATL, needs further empirical validation.

### 3.3 Gender-related regions embedded in category encoding areas

Our study reveals that gender information processing is deeply intertwined with category processing in the brain. Nearly almost regions encoding within-category gender and cross-category gender, can also represent category information. This supports a mixed-selective neural response, aligning with the category-free neural population hypothesis. It has been demonstrated that posterior parietal cortex (PPC) neurons in rat constitute a dynamic network decoded sensory information according to current behavioral demands (Raposo et al., 2014). Consistently, a study focusing on prefrontal cortex (PFC) activity in macaque monkeys also found that the same PFC circuits mediated flexible behavior through dynamical processes that select and integrate task-relevant sensory input (Mante et al., 2013). Linking this category-free neural population account to gender and category processing, our findings suggest that the some brain regions participate in both functions are not rigidly segregated into “gender-specific” or “category-specific” neural subsets. Instead, these regions form a flexible, category-free population, which can be leveraged to extract either category (e.g., distinguish face and body) or gender information (e.g., distinguish male and female). Notably, rMTG’s shared gender representation coexists with its category selectivity, providing robust evidence for mixed-selective pattern of neural responses. Integrating previous findings and our results, the flexible, mixed-selective pattern seems be adopted in many different brain regions, suggesting it may be a general framework for high-level information perception, reflecting a parsimonious neural strategy. In sum, these shed new light on the flexibility and efficiency of neural computations underlying high-level information perception.

### 3.4 Conclusion

The present study provides the first evidence for category-general neural correlates of gender perception across faces, bodies and objects. The results revealed distributed gender processing in occipitotemporal regions, with significant functional connectivity between upstream and downstream areas specifically for body and object gender. More importantly, rMTG supports shared gender representation across categories in a mixed-selective manner and on mid-level abstraction. Moreover, the observed similarity in functional connectivity for face and body gender suggest a shared neural network for processing gender in person-related stimuli. These findings enrich our understanding of the gender representation through local regions and distributed networks, and inspire future on how diverse cues contribute to processing gender or other dimensions.

## 4 Materials and methods

### 4.1 Participants

Twenty-two healthy participants (11 male), aged from 19 to 28 (M ± SD = 22.4 ± 2.8) years, were recruited in the current study. All participants had normal or corrected-to-normal vision, and were naïve to the purpose of the experiments. They provided informed consent before experiments and were paid for their participation. The protocols of the research were approved by the institutional review board of the Institute of Psychology, Chinese Academy of Sciences and adhered to the tenets of the Declaration of Helsinki.

### 4.2 Stimuli

There were six types of images for the main experiment: face, body and object images varying in gender (male and female). Each type of them included 30 different images. Face images were from Chinese male (30) and female (30) movie actors. Body images were the body in underwear below the head and above the knees. Face and body images were downloaded from the Internet. Object images were closely associated with male or female genders, selected from a previous study (Javadi & Wee, 2012).

### 4.3 Procedure

In the fMRI scanner, stimuli were back-projected onto a screen (60 Hz frame rate, 1024 × 768 pixels screen resolution) via a liquid crystal projector and viewed through a mirror mounted on the head coil. The screen was positioned at a distance of 80 cm. Stimuli were displayed using MATLAB (Mathworks, Inc.) together with the Psychophysics Toolbox (Brainard, 1997; Pelli, 1997).

The fMRI experimental runs were arranged in a block design with each run containing 18 blocks. Each run contained 6 conditions of a 2 (gender: male vs. female) × 3 (category: face, body, object) factor design (Fig. 1a), which were presented three times in random order. Each block presented 12 images, where each image with fixation was shown for 0.9 s followed by a 0.1 s fixation between images (interstimulus interval, ISI; except that the first two participants, the presentation durations of each image and the ISI were 0.5 s). A fixation was presented 6 s between blocks. The 12 images included 10 different images and the repetition of two of them (Fig. 1b). All the stimuli were presented on a grey background. The face and body images were cropped into a square with grey background (8.35°×8.35°), and the object images were cropped into a rectangle with grey background (11.12°×8.35°). Participants were instructed to press the button when a consecutive repetition of an identical image was presented (one-back task). Participants completed 6 runs (except that the first two participants completed 7 runs), and each run lasted 5 minutes and 30 seconds. Participants’ mean accuracy on the one-back task was 96.83 ± 3.54%, demonstrating they have focused on the images during scanning.

### 4.4 fMRI scan parameters

Functional and anatomical data were collected using a 3-Tesla Siemens Prisma scanner with 64-channel head coil at the Beijing Magnetic Resonance Imaging Center for Brain Research. Functional data were collected using a T2*-weighted echo planar imaging (EPI) sequence with the following parameters: TR = 2000 ms, TE = 30 ms, flip angle = 70°, FOV = 192 × 192 mm. Volumes consisted of 78 slices (with multiband), with an isotropic voxel size of 2 × 2 × 2 mm. A 3D T1-weighted magnetization-prepared rapid gradient echo (MP-RAGE) image was acquired with the following parameters: 128 sagittal slices, TR = 2600 ms, TE = 3.02 ms, TI = 900 ms, flip angle = 8°, FOV = 256 × 224 mm, isotropic voxel size of 1 × 1 × 1 mm.

### 4.5 fMRI data preprocessing

The fMRI data was processed using SPM12 (http://www.fil.ion.ucl.ac.uk/spm/). The first 3 volumes of experimental run and the first 2 volumes of localizer run were removed to avoid T1 saturation. All functional data was realigned, slice-time corrected, and co-registered with the T1 anatomical image. MVPA and RSA were conducted on unsmoothed data in subject-space. The single-subject accuracy map of searchlight classification in MVPA and the single-subject β weight maps (see details in 2.7.1) of searchlight RSA were normalized to MNI space, and spatially smoothed with a 4 mm Gaussain kernel. For PPI analysis, the anatomical images were spatially normalized to the Montreal Neurological Institute (MNI) template, and normalization parameters were applied to the functional images, and the functional data was also spatially smoothed with a 4 mm Gaussian kernel.

### 4.6 Multivoxel pattern analyses (MVPA)

Data recorded by fMRI from each experimental run was separately modelled with General Linear Models (GLMs), in which each block was modeled as separate regressors in the GLM in addition to 6 head motion parameters. The betas map of 3 times of presentation × 6 (7) runs of each condition from the GLMs were extracted, demeaned to reduce the amplitude effects of different gender, and then submitted to MVPA. The gender MVPA were conducted both within category (face or body or object) and across categories (across face and body, across body and object, across object and body). The category MVPA were conducted on each combination of two categories. We conducted these MVPA using a whole-brain searchlight analyses, with each searchlight consisting 200 voxels. The classification accuracies of each searchlight were assigned to the central voxel, generating whole-brain maps of accuracies. The whole-brain accuracy maps were tested against chance level (50%) using one-sample t-tests to identify regions with above-chance classification, and the results was corrected for the multiple comparisons using a cluster-based Monte Carlo simulation (implemented in the CoSMoMVPA toolbox, (Oosterhof et al., 2016)) at *p* < 0.05. We used a threshold of *p* = .001 at the initial voxel-wise, and 5000 iterations of Monte Carlo simulations (Forman et al., 1995; Goebel et al., 2006). For visualization, maps were projected on a cortex surface of the BrainNet toolbox (Xia et al., 2013).

The within-category MVPA on gender was conducted to explore which brain regions encode the gender of specific category, separately according to the neural pattern evoked by viewing faces, bodies and objects of different gender (i.e., male and female). Leave-one-out cross-validation was implemented with CoSMoMVPA. Specifically, an SVM classifier was trained for gender classification of specific category (e.g., female face versus male face) in all but one run of data, the ‘left-out’ run of data was used to independently test classification of gender within category. This was iterated 6 times (7 times for the first two participants) with each run used once as the left-out test run. The classification accuracy was determined by averaging multiple cross-validation iterations. The overlap of brain regions encoding gender from the three categories were examined to investigate the commonality in their representations.

The cross-category MVPA on gender was conducted to further investigate the brain regions encoding gender across different categories. The analyses were identical to the within-category MVPA on gender, except that the classifier was first trained using the data from one category (e.g., face) and subsequently tested on the data from the other category (e.g., body). For instance, in the classification of gender across face and body, the classifier was trained on face and tested on body, and trained on body and tested on face. The final classification accuracy was determined by averaging the two training and test combinations. A conjunction analysis of the three cross-category MVPA provided more robust evidence for brain regions with shared gender representation.

The MVPA on category was also similar to the within-category MVPA on gender, except that the classifier was trained for category classification (e.g., face versus body). The final map for encoding category information was obtained using a conjunction analysis of category classification between each pairwise categories. The brain regions engaging in category representation were then compared with those brain regions engaging in gender representation to demonstrate the extent of their overlap.

### 4.7 Representational Similarity Analysis (RSA)

To further confirm the regions with shared gender representation, we conducted a RSA using CoSMoMVPA toolbox (Oosterhof et al., 2016). First, for fMRI data, GLM containing predictors of the six conditions and six head motion parameters was conducted in each run, generating SPM_T_ maps for each condition. Neural representational similarity matrix (RSM) was created by calculating the Pearson correlation between the SPM_T_ maps for each pairwise form six conditions. Then the neural RSM was converted into neural RDM using 1-Pearson correlation.

Given that gender representation cannot be completely disentangled from category representation (as revealed by the MVPA results, see 3.2.2 and Fig. 2e), a regression-based RSA was conducted using a whole-brain searchlight analysis (200 voxels per searchlight) to model the neural RDMs as a function of theoretical gender and category RDMs. Theoretical gender and category RDMs (6 × 6 binary matrices) were predictors, with 1 for between-gender/category stimulus comparisons (e.g. male vs. female) and 0 for within-gender/category stimulus comparison (e.g. female vs. female). The neural RDM was fitted via multiple regression (Fig. 1C): 𝑛𝑒𝑢𝑟𝑎𝑙 𝑅𝐷𝑀 ∼ 𝛽_0_ + 𝛽_1_ 𝑔𝑒𝑛𝑑𝑒𝑟𝑅𝐷𝑀 + 𝛽_2_ × 𝑐𝑎𝑡𝑒𝑔𝑜𝑟𝑦 𝑅𝐷𝑀.

Each searchlight’s beta values were assigned to the central voxel, generating whole-brain 𝛽 maps. These 𝛽 maps were entered into one-sample t-tests (against 0), using the identical statistics as for MVPA accuracy (see details in 2.6). The gender-related 𝛽 map was compared with the shared gender-encoding regions revealed by the MVPA to check their consistency.

To further characterize which level of features in the processing hierarchy (low, intermediate, or high-level) drives shared gender representation in rMTG, we submitted the images used in fMRI to a fine-tuned convolutional neural network model (CNN). CNNs have hierarchical architecture matching human visual processing (Eickenberg et al., 2017; Güçlü & van Gerven, 2017). Lower layers capture low-level visual features, while higher layers capture more complex and abstract features (LeCun et al., 2015). Thus, we could infer the hierarchical nature of the share gender representation by identifying the layer whose RDM optimally matches the neural RDM.

Specifically, we fine-tuned a AlexNet pretrained on ImageNet-1K dataset (Olga et al., 2015) to classify gender using backpropagation in PyTorch (Adam et al., 2019). The fine-tuned process was run on a Men/Women Classification dataset (https://www.kaggle.com/datasets/playlist/men-women-classification), using Adaptive

Moment Estimation (learning rate: 0.001, batch size: 32 images, epochs: 10). The dataset contained 2000 training images (1000 males), and 800 validation images (400 males). Training accuracy was 85.0%, validation accuracy was 79.3%. On our images used in fMRI experiment, test accuracies for face and body gender were both 81.7%.

These accuracies are comparable to a prior work showing that AlexNet’s performance of 82.6% on gender classification (Gurram et al., 2024). The test accuracy was 70.0% for object gender. The relatively lower accuracy for object gender discrimination may reflect the inherent ambiguity of gender cues in objects, as object gender is mostly shaped by experience or culture and exhibits greater variability in training images than the biological basis of face and body gender. For instance, some objects, such as a brown comb or a red helmet, are less stereotypical and ambiguous with respective to gender classification.

To construct CNN RDMs, the activation patterns of the fine-tuned AlexNet in response to our experimental images were extracted from five convolutional layers (conv1 to conv5) and two fully connected layers (fc6, fc7). Dissimilarities were calculated using 1-Pearson’s R on activations of each pairwise, generating seven 6× 6 RDMs. We then calculated the neural RDM in the category-general gender-related region (rMTG; MNI: 12, −98, 0; 73 voxels, see Results 3.2.2) for each participant, and correlated participants’ neural RDMs with the CNN RDM from each layer. The correlation coefficients (Pearson r) were Fisher z transformed, and submitted into one-sample t-tests. For comparison, we also computed the correlations between CNN RDMs with neural RDMs in V1 (MNI: 12, −98, 0, 81 voxels) and PFC (MNI: 48, −62, 0, 81 voxels) to illustrate the similarity between CNN layers and neural activities in representative lower and higher brain regions in the processing hierarchy. The hierarchical nature of the neural representation in the three regions (rMTG, V1 and PFC) were elucidated by two linear mixed models fitted to the calculated correlation coefficients across layers. One model included only the linear term ( Correlation ∼𝑙𝑎𝑦𝑒𝑟 + (1|subject) ), while the other included both the linear and quadratic term (Correlation ∼𝑙𝑎𝑦𝑒𝑟 + 𝑙𝑎𝑦𝑒𝑟^2^ + (1|subject)). A quadratic trend of the correlation coefficients across layers supports an intermediate representation in the given brain region, otherwise, it supports a low or high-level representation. We expected a quadratic trend for rMTG and a linear trend for V1 and PFC, according to their neural hierarchy. The optimal layer (with the highest mean correlation across participants) that best matches the neural representation in rMTG was then identified, and further verified by paired t-tests comparing the correlation coefficient at the optimal layer against those at other layers, with FDR correction for multiple comparisons.

To test the generalizability of our findings across different CNN models, we also fine-tuned a pretrained VGG16 for gender classification and selected the five max pooling layers (MP1 to MP5) and the two fully connected layers (fc6, fc7) as the seven layer to compute the RDMs. The whole analysis procedure was identical to that for AlexNet. The fine-tuned VGG16’s training accuracy was 86.0%, validation accuracy was 84.9%. Testing on our images, its test accuracy was 80.0% for face gender, 93.3% for body gender and 71.7% for object gender.

### 4.8 Psychophysiological interaction analyses (PPI)

The PPI analysis was conducted to examine the inter-region functional connectivity for those gender-related clusters and its similarity across categories, further revealing potential functional connectivity shared among the neural representations of gender. Based on the whole-brain searchlight MVPA for gender classification within category, we obtained a brain mask including all brain regions which can decode the gender of faces, bodies and objects. Then, according to the anatomical mask of AAL, the collection mask was divided into 27 clusters, each with more than 50 voxels. All these 27 brain regions were used as seed regions.

The PPI analysis contains the following steps: A GLM was conducted, which contains predictors of the 6 condition (the male and female face, body and object) and 6 head motion parameters with the data concatenated from all the run. For each condition of each participant, the time series (1st eigenvariate) of blood-oxygen-level dependent (BOLD) signal from the seed regions defined above were extracted from this GLM, as physiological variables. The time series were multiplied with the psychological variables (male face vs. female face; male body vs. female body; male object vs. female object) using SPM VOI and PPI modules, as PPI time series. We entered the physiological variable, the psychological variable, the PPI time series and a constant term as regressors to build the PPI GLM models.

For each category, PPI effects between each pair of seed regions were extracted and then performed in a group-level one-sample t-test. The results were corrected by FDR correction at *p* < 0.05 and plotted in brain maps. A significant PPI effects indicated that the gender processing involved functional coupling between the pair of seed regions. Individual 27 × 27 PPI effects matrixes can be obtained for each category. For each participant, we calculated the correlation between each pairwise of two matrixes: *r*_FB_ for the correlation between the matrix of face and body, *r*_BO_ for the correlation between the matrix of body and object, *r*_FO_ for the correlation between the matrix of face and object. All individual correlations were entered into one-sample t-tests. A significant correlation between two PPI effects matrixes indicated that the representation of gender across different categories shared similar functional connectivity.

## Acknowledgement

We thank Amir-Homayoun Javadi for providing the gender-related object stimuli used in this study.

The authors declared no competing financial interest.

## Data availability statement

All data used in the present study could be accessed at https://www.scidb.cn/s/UJFZrm.

## Founding disclosure

This research was supported by grants from the STI2030-Major Projects (2021ZD0203800, 2021ZD0204200), the National Natural Science Foundation of China (32430043, 32500948), Youth Innovation Promotion Association of the Chinese Academy of Sciences, the Key Research and Development Program of Guangdong, China (2023B0303010004), and the Fundamental Research Funds for the Central Universities.

