## Supporting Information for "A shared neural code for gender across faces, bodies, and objects in the human brain"

The RSA results of fine-tuned VGG16 confirmed the findings of fine-tuned AlexNet (Fig. S1). Firstly, all the neural RDMs in the three brain regions are significantly correlated with the CNN RDM in each layer (*ps* < 0.01). Model fitting showed a quadratic trend of the correlation across layers was most robustly found in rMTG (linear $R^2_{adj}$ = 0.86, quadratic $R^2_{adj}$ = 0.95; linear BIC = -71.67, quadratic BIC = -211.98), whereas the linear and the quadratic trend fit comparably well in V1 (two models' $R^2_{adj}$ were 0.84; linear BIC = -150.76, quadratic BIC = -145.73) and PFC ( two models' $R^2_{adj}$ were 0.91; linear BIC = -168.81, quadratic BIC = -164.69) and PFC. In rMTG, the strongest correlation was found at layer 4 (i.e., MP4), significantly higher than those at other layers (*ps* < 0.01, see Fig. S1 and Table S1).

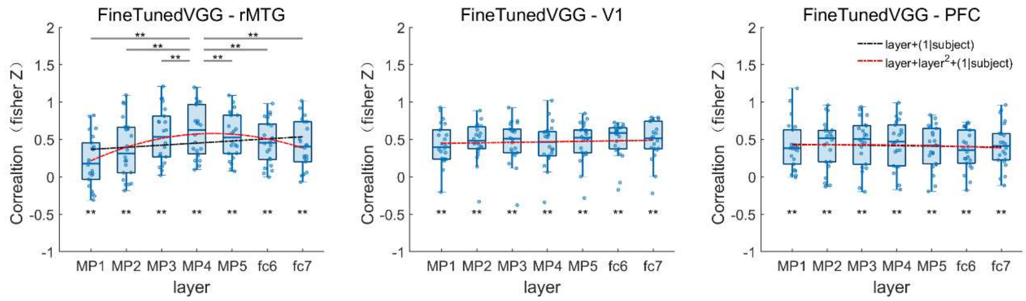


Figure S1. RSA results linking rMTG (left panel), V1(middle panel) and PFC (right panel) to fine-tuned VGG16 layers. * $p < 0.05$, ** $p < 0.01$ (FDR- corrected). The data predicted by the linear and quadratic model were plotted using red and black dashed lines.

Table S1 Statistical details for the RSA results linking the representation of brain and CNN

| Layer | Alexnet | VGG |
|---|---|---|

| | MTG | V1 | PFC | MTG | V1 | PFC |
|---|---|---|---|---|---|---|
| Layer 1 | 0.15 ± 0.29 | 0.35 ± 0.18 | 0.28 ± 0.23 | 0.21 ± 0.34 | 0.44 ± 0.27 | 0.41 ± 0.32 |
| Layer 2 | 0.34 ± 0.36 | 0.42 ± 0.21 | 0.40 ± 0.32 | 0.37 ± 0.39 | 0.48 ± 0.26 | 0.43 ± 0.33 |
| Layer 3 | 0.68 ± 0.33 | 0.33 ± 0.36 | 0.41 ± 0.36 | 0.55 ± 0.36 | 0.47 ± 0.28 | 0.46 ± 0.32 |
| Layer 4 | 0.70 ± 0.32 | 0.34 ± 0.38 | 0.42 ± 0.35 | 0.60 ± 0.34 | 0.44 ± 0.28 | 0.45 ± 0.32 |
| Layer 5 | 0.53± 0.43 | 0.44 ± 0.28 | 0.48 ± 0.37 | 0.55 ± 0.31 | 0.47 ± 0.27 | 0.40 ± 0.28 |
| Layer 6 | 0.58 ± 0.34 | 0.45 ± 0.30 | 0.45 ± 0.33 | 0.46 ± 0.29 | 0.49 ± 0.24 | 0.36 ± 0.27 |
| Layer 7 | 0.55 ± 0.35 | 0.46 ± 0.29 | 0.46 ± 0.33 | 0.42 ± 0.32 | 0.49 ± 0.26 | 0.42 ± 0.30 |

Note: Values are Pearson correlations (Fisher z transformed) averaged across participants (mean ± SD).
